# Invasion and intracellular proliferation of non-pathogenic *E. coli* in model infection systems

**DOI:** 10.1101/2025.08.12.669981

**Authors:** Lachlan Chisholm, Rebecca von Essen, Alaska Pokhrel, Bill Söderström

## Abstract

Uropathogens are typically distinguished from commensal or laboratory *E. coli* strains by defined pathogenic behaviours. In this study, we compared the behaviour of non-pathogenic *Escherichia coli* to that of uropathogenic *E. coli* in diverse infection models at a single-cell level using live-cell fluorescence microscopy. Using *in vitro* microfluidics and *ex vivo* mouse urinary tract infection model systems, we demonstrated that the standard laboratory *E. coli* strain MG1655 can behave like a pathogen, invading and proliferating inside human and mouse urothelial cells. Upon exposure to stress by the addition of urine, we observed that MG1655 undergoes infection-related filamentation, and then subsequent reversal back to rod shape when the stress is removed. This observation supports that infection-related filamentation should be regarded as a stress response rather than a virulence factor. Taken together, our results show that under infection-favourable circumstances, even a well-studied non-pathogenic laboratory *E. coli* strain can exhibit what are considered pathogenic traits.

## Introduction

For sustained survival, bacteria must overcome significant challenges within different host niches, including hostile environmental conditions with limited nutrient availability (1). Particularly suited for this challenge, the Gram-negative, rod-shaped *Escherichia coli* is one of the most genetically heterogeneous bacterial species. In a human context, it ranges from gut-residing commensal strains to several different pathotypes, which colonise a variety of different niches, including the bladder (2). Pathotypes of *E. coli* are broadly separated into the intestinal pathogenic *E. coli* (InPEC) and the extraintestinal pathogenic *E. coli* (ExPEC) (3), which are further subdivided, depending on their specific pathogenic traits and site of infection.

ExPEC express virulence factors, such as adhesive surface structures (*e.g.*, type 1 fimbriae and P fimbriae), siderophores, and toxins that confer greater fitness in diverse host microenvironments. These virulence factors support attachment, invasion, and nutrient scavenging for successful survival and colonisation of the host (4–7). Commensals that colonise the gastrointestinal tract, however, also express these virulence factors (8). In particular, type 1 fimbriae and the tip adhesin FimH, are expressed by both commensal and most pathogenic *E. coli*, mediating adhesion to host epithelial cells (4, 9). Type 1 fimbriae are also expressed by uropathogenic *E. coli* (UPEC) and are necessary for urothelial cell binding and invasion (10). Although UPEC express a FimH variant with a greater ability to bind glycoproteins found on the bladder cell surface (11), both commensal and pathogenic strains of *E. coli* produce this adhesion complex. This raises the question of how commensal *E. coli* behave if they become intracellular?

Urinary tract infections (UTI), caused primarily by UPEC, are among the most common detrimental bacterial infections, affecting approximately 400 million people in 2019 (12, 13). UPEC are a genetically diverse pathotype of the ExPEC group, which, unlike other *E. coli* pathotypes, have no consistent set of conserved virulence genes (14). The key first steps in the commonly accepted model of UPEC pathogenesis are bacterial invasion and replication into intracellular bacterial communities (IBCs) (15, 16). IBC formation allows UPEC to evade host defences, leading to maturation and eventual dispersal from the host cell as rod-shaped or filamentous bacteria (13). Filamentation is the inhibition of cell division while DNA replication and bacterial cell elongation continue (17). The SOS DNA repair pathway is responsible for regulating filamentation during normal vegetative growth under unfavourable conditions. This pathway is activated in response to external stressors, such as antibiotics and UV radiation, as a survival mechanism against DNA damage (18–20). However, infection-related filamentation during UTI has been demonstrated in bacteria that lack SOS response-related genes (e.g., *sulA* and *ymfM*) (21). While the exact molecular cues leading to filamentation have not been exhaustively defined, it is known that the cell division-related SPOR-domain protein DamX is involved in the regulation of the filamentous phenotype (22). It is generally believed that the filamentous form helps UPEC spread, reaching neighbouring bladder cells to infect, and is regarded as a virulence factor (23). In the UTI context, urine is known to induce the filamentation response (21, 22, 24, 25), suggesting that specific components (such as small-molecule constituents (25)) of urine are involved in the deactivation and regulation of the division machinery.

The understanding of UPEC virulence has been largely based on model UPEC strains in model infections (26, 27). Studies evaluating the behaviour of non-pathogenic clinical *E. coli* isolates and laboratory strains have yielded conflicting results regarding invasion, intracellular survival, or filamentation (28–30). In mice, non-pathogenic laboratory K-12 strains invade surface bladder urothelial cells to a limited capacity. However, these strains were unable to survive intracellularly and did not develop into large IBCs or persist within mouse bladders (28). By contrast, in human 3D-microtissue models, commensal *E. coli* isolates from the urine of healthy volunteers underwent the UPEC invasion cycle and adopted a filamentous morphology in late-stage invasion (30). The non-pathogenic laboratory strain MG1655, however, did not filament during infection in this model (30).

Mapping how *E. coli* behaves at the single-cell level during invasion and infection will elucidate whether virulence-associated traits are ‘reserved’ for pathogenic strains or are a broadly shared stress response across the species. Fundamental questions remain regarding whether domesticated non-pathogenic laboratory strains (*e.g.,* MG1655) of *E. coli* behave similarly to UPEC strains with respect to invasion, IBC proliferation, or filamentation in urothelial cells. To shed light on these questions, we used diverse infection models, including *in vitro* petri dish and microfluidic flow models, and an *ex vivo* mouse bladder sheet model, in combination with live-cell fluorescence microscopy to examine bacterial behaviour at single-cell resolution.

We examined the invasion and intracellular proliferation of the standard non-pathogenic laboratory *E. coli* strain MG1655 in comparison to a frequently used cystitis UPEC strain, UTI89. We found that, under our laboratory conditions, the non-pathogenic laboratory strain behaved similarly to the pathogenic UPEC strain during the infection cycle. These results clearly show that commonly used laboratory strains of *E. coli* can effectively invade and proliferate inside human and murine host cells and suggests that morphological changes to filamentous forms should be regarded as a survival stress response rather than a virulence factor.

## Results

### Non-pathogenic E. coli laboratory strain MG1655 invade, proliferate, and form intracellular bacterial communities (IBCs) inside cultured host cells

UPEC invasion and morphological behaviour in host cells in various UTI models is rather well understood (31). In this study, we used a human urothelial cell line PD07i (32) grown as a monolayer to model the outermost layer of the bladder environment in a simple way. Importantly, the PD07i cells retain key aspects of differentiated bladder umbrella cell tissue, including cytokeratin and uroplakin expression, important for maintaining cellular structure and a key receptor for the bacterial invasion cascade (33). In our experimental setup, the bladder cells expressed both E-cadherin and ZO-1 (also called tight junction protein 1, TJP1), indicating the formation of a tight cell barrier reminiscent of that of bladder tissue (Fig. 1A-B) similar to what has been demonstrated in previous reports (34).

**Figure 1.**
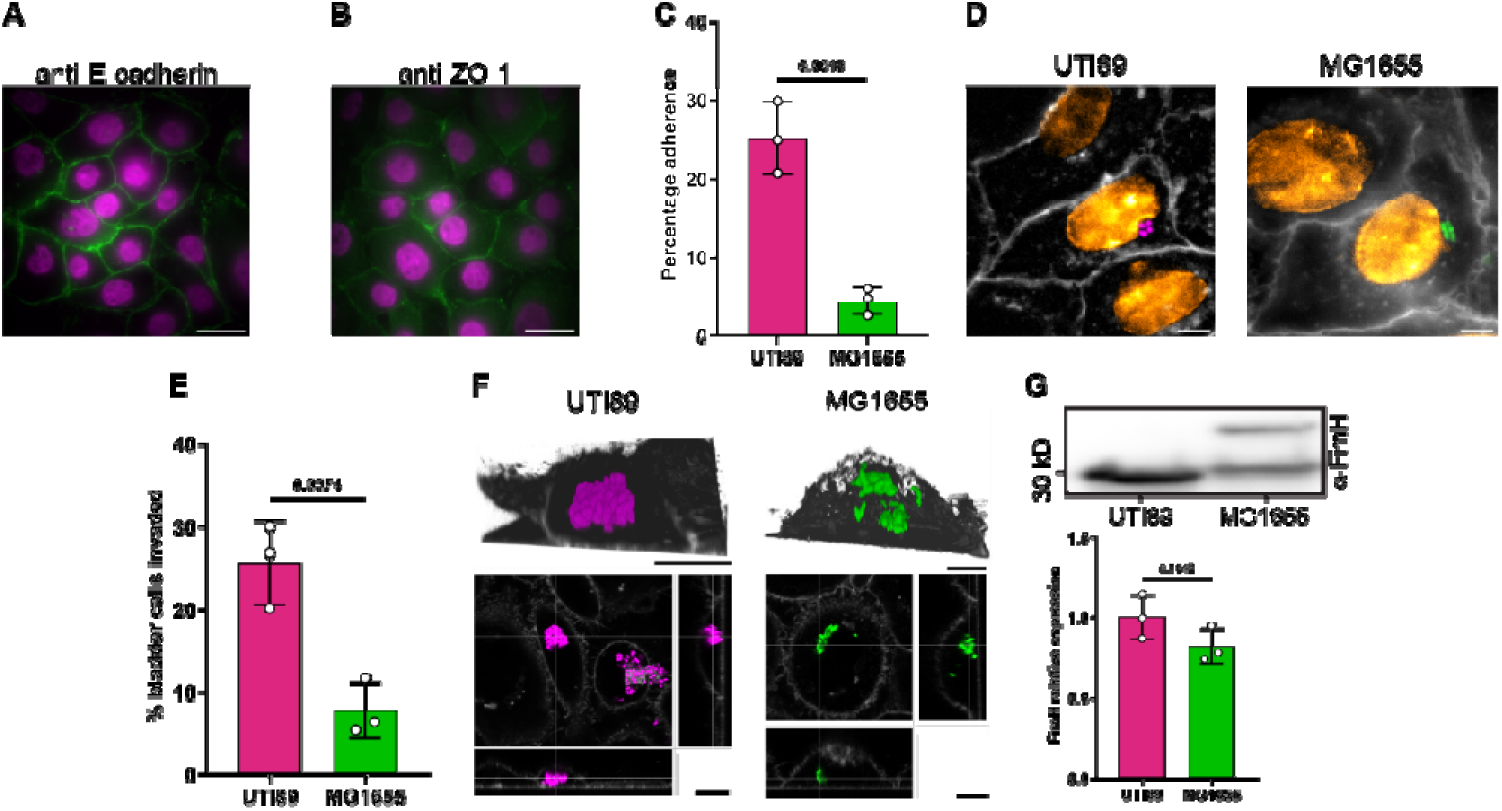
Invasion and IBC formation of various E. coli strains in model infections. Confluent monolayers of PD07i cells stained for antibodies against (**A**) E-cadherin or (**B**) ZO-1 (green). Nuclei stained with NucSpot Live 650 (magenta). **C**, Initial adhesion of UTI89 and MG1655 onto bladder cells relative to the amount of bacteria in the initial inoculum. n = 3. **D**, Live cells in dishes were challenged with either UTI89 or MG1655 E. coli strains. Representative images of bacterial invasion of UTI89 (magenta) and MG1655 (green) highlighting invasion within one host cell in the petri dish model. Host cell membrane stain with CellBrite Steady 405 Membrane stain (pseudo-coloured gray). Nuclei stained with SYTO Deep Red Nucleic Acid Stain (pseudo-coloured hot orange). **E**, Average invasion of urothelial cells by respective strain, n = 3. **F,** 3D-renderings of representative confocal Z-stacks and corresponding cross-sectional views showing IBC formation of UTI89 and MG1655 within host cells. **G**, Quantitative immunoblot showing production levels of FimH in the bacterial strains prior to infection. Graph displaying FimH protein expression of the lower MG1655 band normalised to the average FimH production of UTI89. n = 3. Scale bars in A and B = 20 µm, D = 5 µm and F = 10 µm. Data in **C**, **E** and **G** plotted as mean ± SD. p-values were determined by an unpaired t-test.

To evaluate whether the non-pathogenic *E. coli* strain MG1655, lacking O-antigens, behaves like a prototypical clinical uropathogenic *E. coli* clinical isolate, UTI89, at key stages of the infection, we initially assessed the ability of both bacterial strains to attach to the surface of the bladder cells in a semi-static petri dish infection model. We found that MG1665 remained significantly less adhered to bladder cells compared to UTI89 relative to the initial inoculum (4.53 ± 1.80 % vs 25.22 ± 4.55 %, respectively, mean ± SD, n = 3) (Fig. 1C), consistent with previous trends in published data (35). We then challenged PD07i cells with UPEC strain UTI89, constitutively expressing msfGFP for visualisation (25), examining the invasion frequencies 21 hours post-infection (36). UTI89 invaded urothelial cells with a frequency of 25.63 ± 5.11% (Mean ± S.D., n = 346) (Fig. 1D-E), comparable to previous published data (37).

Next, using fluorescence microscopy, we determined to what extent non-pathogenic *E. coli* would also invade PD07i cells and form IBCs. The formation of IBCs is a hallmark of pathogenic strains during the intracellular phase of the infection cycle in the urinary tract (38). Imaging after the initial 6 hours post-infection clearly showed that MG1655 successfully invaded the urothelial cells (Fig. 1D). After 21 hours post-infection, however, we found that the non-pathogenic strain invaded at significantly lower frequency compared to UTI89, 7.78 ± 3.45 % (Mean ± S.D., n = 411) (Fig. 1E). Following the overnight incubation, confocal imaging further revealed that both strains formed IBCs inside the host cells, however, IBCs formed by the non-pathogenic strain appeared to be less dense, potentially due to lower levels of FimH expression (Fig. 1F). This is consistent with previous studies having shown that IBC formation is dependent on FimH expression (30).

Given the difference in adhesion, invasion frequencies and the appearance of the IBCs between the two strains, we wondered if this possibly could be due to differential production levels of fimbriae, the cellular structures important for bacterial adhesion and invasion (32, 39). *E. coli* encodes 12 different types of fimbriae, of which the best characterised is type-1 fimbriae capped by the adhesin FimH, which is known to bind to mannose residues on the bladder cell surface receptors such as Uroplakin 1a (40, 41). We probed FimH levels prior to bacterial infection of bladder cells by quantitative immunoblotting, observing that relative to the cellular levels of FimH in UTI89, MG1655 produced less FimH (0.82 ± 0.11, mean ± SD, n = 3) (Fig. 1G). For the non-pathogenic strain, there was also a higher molecular weight band present of unknown origin (Fig. 1G). This difference in FimH production may be attributable to the significantly less adhesion and invasion of the non-pathogenic strain compared to the pathogenic strain.

### Non-pathogenic laboratory strains undergo infection-related filamentation similar to model pathogens

After IBC maturation, a key stage in the uropathogenic *E. coli* morphology cycle during urinary tract infection is filamentation (42). Where the bacterial cells arrest the cell division machinery but continue to grow in length, resulting in cells with body lengths exceeding several hundred micrometres (24). This phenomenon has been observed in various *in vitro* and *in vivo* mouse models, as well as in clinical samples from patients with active UTI episodes (23, 24, 28, 38).

Eventually, this leads to host cell rupture and dispersal of UPEC into the host lumen and has been implicated as a persistence mechanism against host immune responses (43). To evaluate whether the filamentation response is specific to pathogens or a broadly adopted response, bladder cells were again challenged with either MG1655 or UTI89. Here, we used both the semi-static dish model and a microfluidics-based *in vitro* infection model with continuous flow of media and nutrients, as this has previously been shown to be beneficial for filament formation (24, 25, 44). For the flow model, urothelial cells seeded in IBIDI microfluidic flow cells were challenged with bacteria under a constant flow of 15 µL min^-1^. This flow rate creates hydrodynamic shear rates (4.9 s^-1^) based on chamber dimensions and a volumetric flow rate that are similar, or slightly higher, to rates that urothelial cells experience *in vivo* (24), promoting conditions to induce infection-related filamentation (44). After 6 hours post-infection, the culture media was supplemented with 50 µg mL^-1^ gentamycin to eliminate extracellular bacteria (25), and the infection was allowed to progress for an additional 18 – 20 hours to generate IBCs. Following this incubation period, the culture media was exchanged for human urine in the flow system. To maximise the likelihood of filamentation, only urine with a urine specific gravity (USG) between 1.025 and 1.030 g mL^-1^ and a pH between 5 and 5.5 was used, as these parameters have previously been shown to be critical to induce filamentation (25, 44). Following another ∼ 18 hours of incubation, the resulting samples were collected from the outlet of the flow channel, placed on a premade agarose pad and directly imaged. To distinguish between bacterial cells slightly elongated due to stress by exposure only to urine, and filamentation specifically due to intracellular host conditions, an ‘infection-related filament’ was defined as any bacterial cell that was ≥10 µm in length, which is >2.5 times the average length of *E. coli* grown in LB cultures (44).

Expectedly, whilst the urine dish model did elicit a slight filamentous response in UTI89 (Ave_length_ = 9.48 ± 10.91 µm (n = 4200, mean ± SD), Fig. 2A), the bacterial cells extracted from the urine flow infection showed a significantly stronger filamentation response, with the average cell lengths of 20.88 ± 32.38 µm (n = 5002, mean ± SD, Fig. 2A). Similarly, MG1655 cells extracted from urine flow infections elicited a significantly stronger filamentation response than cells extracted from the urine dish infections, 47.53 ± 59.67 µm (n = 1537, mean ± SD) compared to 21.94 ± 28.16 µm (n = 2257, mean ± SD) from the urine dish model (Fig. 2A). We observed that flow infections without urine did not induce filamentation in either UTI89 or MG1655 (2.159 ± 0.5480 µm, n = 6404 and 3.644 ± 1.407 µm, n = 3439, mean ± SD, from three independent experiments respectively (Fig. 2A). No filamentation was observed for bacteria grown statically overnight in urine only (Supplementary Fig. S1).

**Figure 2.**
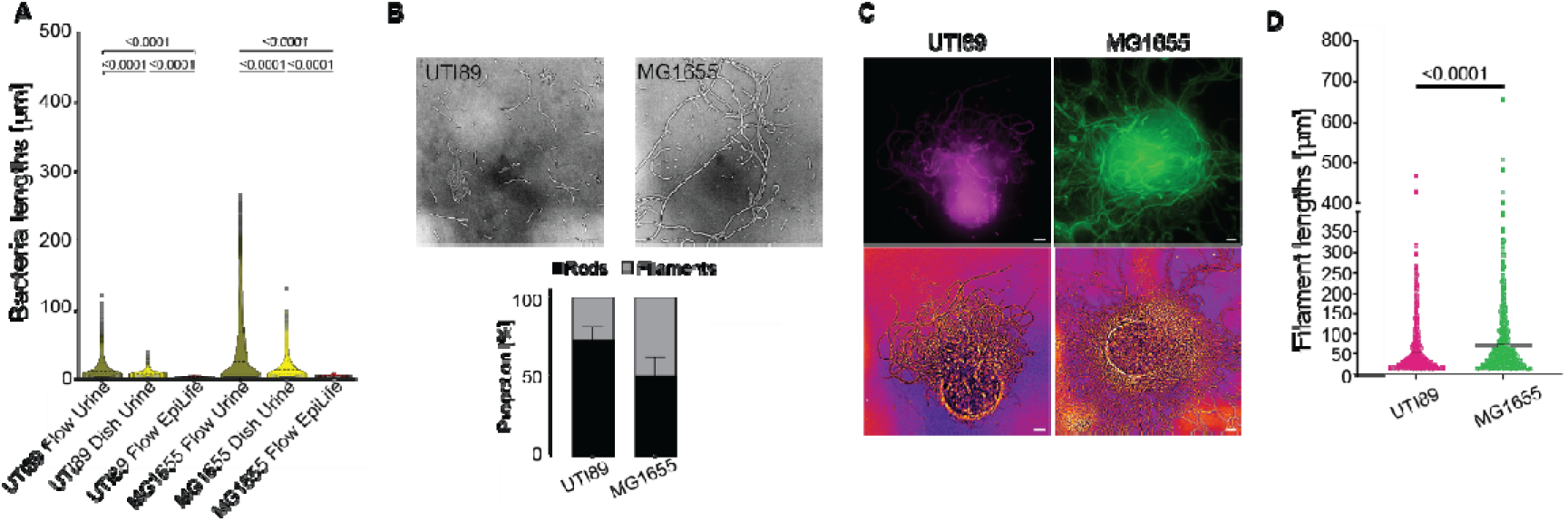
Filamentation of non-pathogenic E. coli in model microfluidics flow infection. **A**, Length distribution of UTI89 and MG1655 extracted from infections with urine flow, semi-static infection with urine and EpiLife flow. n _≥_ 1537 cells of each strain per infection. p-values were determined by one-way ANOVA comparing strain-specific infection conditions. **B**, Representative bright-field images of the different bacterial cell phenotypes extracted from supernatant after 18 hours of urine flow (46 hours post-infection). Bottom middle: ratio of filaments to rods in the supernatant of samples (mean + SD, n = 3 independent urine flow infections with each strain). **C**, Representative images of filamentation of each strain as bacteria effluxed from urothelial cells in the flow model 46 hours post-infection. Both bacterial strains produced cytoplasmic GFP from plasmid pGI5, pseudo-coloured to visually distinguish strains (UTI89:magenta, MG1655:green). Bright-field images are displayed using the FIJI fire LUT. **D**, Length distribution of filaments from each strain extracted from the flow model 46 hours post-infection (n _≥_ 305 filaments from each strain). UTI89 = 54.26 ± 56.81 µm and MG1655 = 73.55 ± 81.06 µm (mean ± S.D., n = 810 and 579 respectively). p-values determined by an unpaired t-test. Scale bars B and C = 10 µm.

As the urine flow infections resulted in the strongest filamentation response, we conducted follow-up infections to observe if there were morphological differences in the filamentation response between the two strains during this model. Consistent with previous studies (24, 25, 44), UTI89 exhibited a mixture of filaments and rods, with the average filament length measuring 54.26 ± 56.81 µm (mean ± SD, n = 810) (Fig. 2B-D). The percentage of filaments in the UTI89 infections was 27.13 ± 8.65 % (mean + SD, n = 4580) of the total cell population (Fig. 2B). Similarly to UTI89, MG1655 also exhibited a mixture of filaments and rods, however filamentous MG1655 cells constituted almost half of all cells, 49.99 ± 11.98 % (mean ± SD, n = 1755) (Fig. 2B). Filamentous MG1655 cells were on average significantly longer than the UTI89 filaments (Fig. 2C - D), with average lengths MG1655_Lengths_ = 73.55 ± 81.06 µm (mean ± SD, n = 579) (Fig. 2D).

Our observation that the non-pathogenic laboratory strain underwent infection-related filamentation was unexpected, as previous studies have shown that MG1655 did not undergo infection-related filamentation, however, the urine parameters for filamentation were not specifically determined in those studies (28, 30).

### MG1655 largely follows the morphology patterns of UTI89 in an ex vivo bladder sheet mouse model

A previous study indicated that non-pathogenic and pathogenic *E. coli* invaded bladders to a similar extent *in vivo*, however, the non-pathogenic bacteria did not persist, multiply intracellularly, or exit host cells based on bacterial enumeration CFU counts (28). Given that the non-pathogenic strain both invaded host cells and formed filaments in our models, we wanted to determine whether the non-pathogenic strain behaved similarly in an *ex vivo* mouse bladder sheet model. We used an *ex vivo* mouse urothelial sheet model, which yields a comparable infection burden to *in vivo* mouse models (45). In this model, the urothelium and lamina propria are separated from the muscle layer, resulting in a thin urothelial sheet. One noticeable difference from previous *in vivo* studies is that the bladder sheets lack a fully intact immune system, however, immunoactivity is maintained with several bladder-resident immune cells remaining active (45).

We then isolated urothelial sheets from naïve female C57Bl/6 mice and challenged them with UTI89 or MG1655. After overnight incubation, sheets were stained for nuclei before being splayed on agarose pads and imaged. Both strains invaded host epithelial cells and proliferated intracellularly (Fig. 3A), supporting the notion that non-pathogenic *E. coli* invade the urothelium in mice. However, while clearly invasive, MG1655 invaded significantly fewer cells compared to UTI89 (Fig. 3B). A noticeable difference was that UTI89 more readily formed dense IBCs, while MG1655 was more diffused throughout the host cytoplasm (Fig. 3C). This correlated with lower FimH production in MG1655 compared to UTI89 (Fig. 1G).

**Figure 3.**
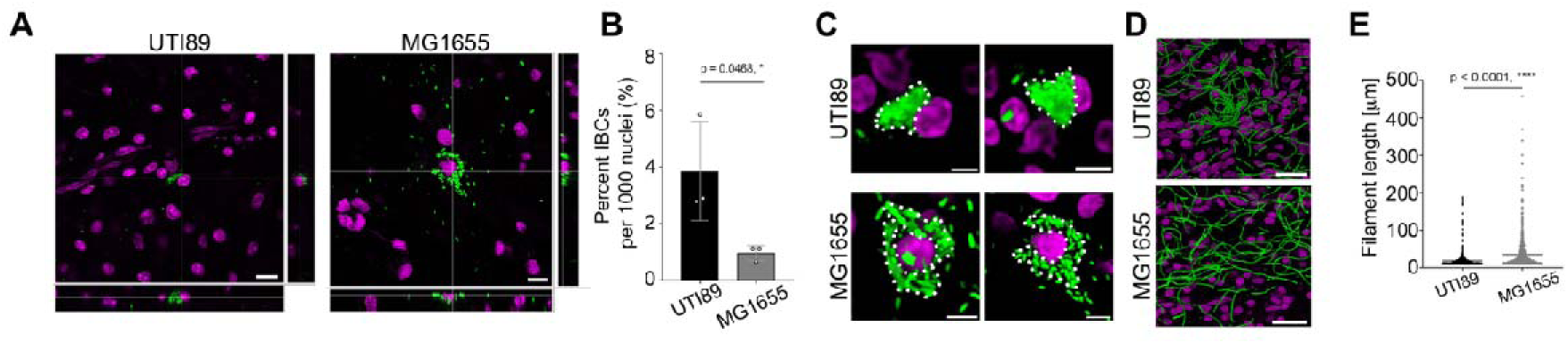
The non-pathogenic E. coli strain MG1655 behaves like a pathogen during infection of ex vivo bladder sheets. Peeled urothelial sheets from C57Bl/6 mice were infected with pathogenic UTI89 or non-pathogenic MG1655 for 4 hours, treated with gentamycin and incubated overnight. Sheets were stained with NucSpot Live 650 (magenta) to visualise nuclei by confocal microscopy. **A,** Representative cross-sectional views highlighting intracellular IBC formation of both UTI89 and MG1655 in bladder sheets. **B**, Graph shows quantification of A, invasion frequencies for the two E. coli strains. **C**, Images show close-up views of representative infections in A. **D**, Images show urine-exposed sheets and bacterial filaments visualised by confocal microscopy. **E**, Length distribution of UTI89 and MG1655 filaments produced inside urothelial sheets. All p-values were determined by an unpaired t-test. Scale bars A, C = 10 µm, D = 50 µm.

Next, to test whether MG1655 would undergo infection-related filamentation similar to UTI89 in *ex vivo* bladder sheets, we exposed infected sheets to human urine, with appropriate parameters for filamentation (*i.e.,* pH 5.5 and USG 1.025 (25)) overnight. UTI89 filamented (Fig. 3D) as expected and previously reported (45), and to our surprise, the non-pathogenic strain exhibited extensive filamentation (Fig. 3D). Noticeably, the average length of the UTI89 filaments were significantly shorter than the MG1655 filaments in both models, with UTI89 lengths Ave*_ex vivo_* = 19.11 ± 18.05 µm (n = 734, Fig 3E) (Ave*_in vitro_* = 54.26 ± 56.81 µm, Fig 2D), and MG1655 Ave*_ex vivo_* = 35.38 ± 41.59 µm) (n = 810, Fig 3E) (Ave*_in vitro_* = 73.55 ± 81.06 µm, Fig 2D).

### Filament reversal dynamics of MG1655 mirror that of UTI89

The ability of filamentous UPEC to survive and initiate secondary infection is dependent on efficient reversion to viable rod-shaped bacteria, thereby completing the dispersal phase (19, 46). We therefore wanted to determine whether filaments of non-pathogenic MG1655 would efficiently revert to rods when urine stress was removed. We collected the supernatant from the urine-exposed bladder sheets, centrifuged, and the resuspended pellet was placed on agarose pads. The pads were placed under a 100x objective in a 37 °C chamber and imaged every 10 minutes for at least 120 minutes. In this period, we observed that the filaments either elongated, divided and reverted to rods or did not divide and lysed (44). A majority of UTI89 filaments reverted back to rods (∼83%, n_tot_= 109) (Fig 4A) by 120 minutes, whereas a larger number (∼34%, n_tot_ = 151) of MG1655 filaments did not grow or exploded during imaging (Fig. 4A-B).

**Figure 4.**
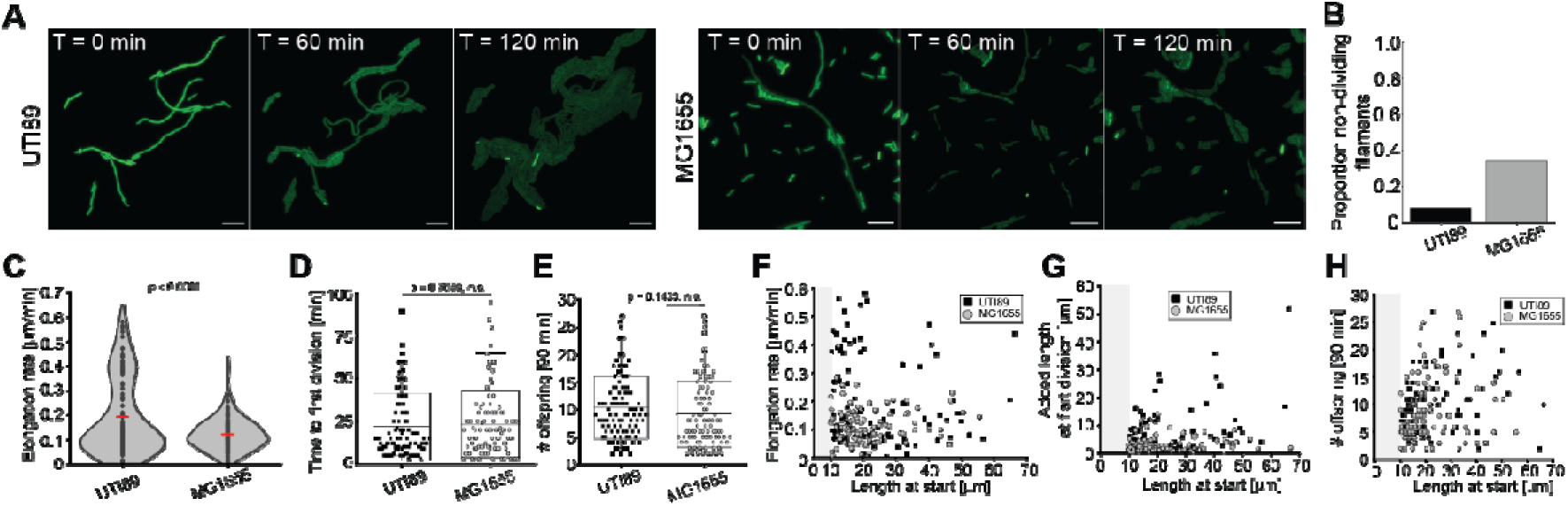
Infection-related filaments reverse over time after exposure to urine. **A**, Samples were collected from the supernatant of the experiments, placed on agarose pads and directly imaged. Snapshots of filament reversal of UTI89 and MG1655 over time. Scale bars = 10 µm. **B**, Proportion of lysed or non-dividing filaments of each strain. **C**, Elongation rates for filaments placed on agarose pads after exposure to urine. Red lines indicate mean. **D**, Time to first division for each strain. T_UTI89_ = 22.13 ± 19.46 min (n = 98), T_MG1655_ = 23.17 ± 19.27 min (n = 96). **E**, Graph shows the number of offspring after 90 min of imaging. Offspring_UTI89_ = 10.4 ± 5.6 (n = 98), offspring_MG1655_ = 9.2 ± 6.1 (n = 96). **F**, Bacterial cell length at the start of imaging versus elongation rate. UTI89: R^2^ = 0.012, Pearson’s r = -0.11. MG1655: R^2^ = 8E^-4^, Pearson’s r = 0.028. **G**, Bacterial cell length at the start of imaging versus length added up to the first division during imaging. UTI89: R^2^ = 0.14, Pearson’s r = 0.38. MG1655: R^2^ = 0.067, Pearson’s r = 0.082. **H**, Bacterial cell length at the start of imaging versus number of offspring after 90 min. UTI89: R^2^ = 0.097, Pearson’s r = 0.31. MG1655: R^2^ = 0.015, Pearson’s r = 0.12. Boxes in plots in **D** and **E** represent SD, with midline showing averages. Whiskers indicate 95% of the data. p-values in **C**, **D**, and **E** were determined by unpaired t-tests.

The observed elongation of the filaments before the initiation of division correlates with a recovery phase after stress, when a nutrient-rich environment is supplied after the nutrient-poor urine (47). Quantification of filament elongation for the two strains indicated a higher rates for UTI89 (rate_UTI89_= 0.191 ± 0.15 µm min^-1^ (n = 100), rate_MG1655_ = 0.121 ± 0.06 µm min^-1^ (n = 96), Fig. 4C), however no significant difference was observed for the average time to the first daughter cell division (T_UTI89_ = 22.13 ± 19.46 min (n = 98), T_MG1655_ = 23.17 ± 19.27 min (n = 96)) (Fig 4D), nor in the number of offspring after 90 minutes of imaging (offspring_UTI89_ = 10.4 ± 5.6 (n = 98), offspring_MG1655_ = 9.2 ± 6.1 (n = 96)) (Fig. 4E). These results were comparable what has been observed previously for filament reversal in an *in vitro* UTI flow model (44).

Further analysis between the initial length of the filaments vs elongation rate, added length at first division or offspring after 90 min showed that these parameters were not correlated (Fig. 4F-H). These findings suggest that although the filaments of the non-pathogenic strain may not be as adept at surviving in urine as the pathogenic UTI89 strain, nor as efficient in switching from nutrient-poor to nutrient-rich environments, MG1655 filaments can behave similarly to the pathogenic strain when provided the right conditions.

## Discussion

Successful bacterial colonisation and persistence within niches are key survival challenges bacteria face within hosts (48). One strategy that *E. coli* uses is adherence to epithelial cells with subsequent invasion, intracellular proliferation, and survival (49). The expression of surface adhesion structures, like the type 1 fimbriae, are necessary for attachment and invasion into host cells (50). For intracellular pathogens, such as uropathogenic *E. coli,* invasion into bladder epithelial cells leads to intracellular bacterial communities, providing protection from host immune responses and antibiotic exposure (4). It is not well characterised, however, if non-pathogenic *E. coli* adopts a similar approach.

In the current study, we investigated the initial adhesion, invasion, intracellular survival, and morphology changes of pathogenic and non-pathogenic model *E. coli* strains side-by-side. Our observations of how the common non-pathogenic laboratory strain MG1655 behaves similarly to the model UTI strain UTI89 raises questions regarding whether survival tactics overlap with virulence phenotypes, when key survival strategies are shared between non-pathogenic and pathogenic *E. coli*.

Results from *in vitro* and *ex vivo* infection models clearly show that MG1655 can mirror the behaviour of UTI89 at the single-cell level, with both strains attaching to and invading urothelial cells. Similar results are observed for commensal strains isolated from healthy donors (30, 37), supporting that invasion into urothelial cells is a broadly shared strategy rather than a specific virulence trait.

Once inside the host, both strains formed IBCs. However, in the *in vitro* dish and *ex vivo* mouse bladder sheets, IBCs were generally not as dense in MG1655 infections compared to UTI89, which may be due to lower expression of FimH. This is consistent with data linking FimH to dense IBC formation in a human microtissue bladder model (30).

Our data showed that MG1655 underwent infection-related filamentation in both cultured cells and mouse urothelial sheets. This was surprising because filamentation is thought to be a virulence mechanism activated by the innate immune system (18, 51), and MG1655 reportedly does not filament in similar settings (30). A key difference between the studies is that previous data were based on pooled urine, whereas the current study used urine from single donors. Pooled human urine has been shown to artificially influence factors within urine, such as nutrient levels and host proteins, which may not accurately reflect individual patient samples, especially as infection-related filamentation has been shown to be dependent on factors such as pH and specific gravity (25, 52, 53). We also observed that the strongest filamentation response was due to exposure to urine flow and not attributable to media changes or urine alone. Filamentation as a stress response is normally regulated by the SOS pathways (18, 20). However, deletion of SOS pathway genes *sulA* and *ymfM* did not attenuate filamentation in UPEC during infection, suggesting a different activation pathway for the infection-related filamentation stress response (21, 24).

Our data shows that the non-pathogenic *E. coli* laboratory strain MG1655 also filamented during infection, suggesting that infection-related filamentation is a stress-related survival mechanism rather than a virulence factor. If the conditions are right, even laboratory strains can inhibit cell division, resulting in infection-related filamentation upon exposure to urine. Overall, MG1655 exhibited the strongest filamentation response, with an average length ∼ 20 % longer than that of the UTI89 strain. While the MG1655 filaments were longer, they were not equally successful in their reversal to rods compared to the UTI89 strain, with some 40 % dead or dying from bursting during imaging.

Why MG1655 cells proliferate intracellularly and subsequently undergo infection-related filamentation in the *ex vivo* model, while they do not survive in an *in vivo* infection, is not fully understood. One explanation is that in a live mouse, additional host immune factors are present that are not found in the *ex vivo* sheets. More generally, important differences between various models of UTI and other infection studies include human or mouse origin and sex, presence or absence of immune cells, controlled composition of urine parameters and inclusion of flow (54). These rather complex differences in model parameters are bound to influence the outcome of the infection results, which is likely a reason as to why non-pathogenic laboratory strains of bacteria have not previously been observed surviving, proliferating, or filamenting in intercellular host cells at the organism level.

In summary, we propose that, if given the opportunity, any laboratory strain *can* behave similarly to their pathogenic cousins. As non-pathogenic bacteria can be used in less restrictive biosafety level laboratories (*e.g.,* BSL1), we believe this could allow diverse laboratories to conduct “infection-type experiments” without the need for extensive upgrades to facilities. This also opens the floor for discussion on what factors drive disease and, on a fundamental level, what it means to be classified as a pathogen.

## Author contributions

B.S and L.C. conceived the study. All authors performed experiments. R.v.E., L.C. and B.S. analysed the data. B.S. and R.v.E. acquired funding. L.C. and B.S. drafted the manuscript with editorial input from all authors. All authors approved the final version of the manuscript.

## Acknowledgments

The authors wish to thank Fiona Ryan, manager at the ERNST facility at UTS, for the kind support in continually providing mouse bladders. The authors warmly thank Dr Molly Ingersoll for critical reading and critique of the manuscript. The authors acknowledge the use of the Leica Stellaris 8 confocal microscope in the Microbial Imaging Facility at the Australian Institute for Microbiology & Infection in the Faculty of Science at UTS. BS is supported by the Australian Research Council through a Future Fellowship (FT230100062). RvE acknowledge financial support from The Petersenska Hemmet Foundation, The Carl Erik Levin Foundation, The Löfvenskjöld Travel Grant Fund administered by the House of Nobility, Sweden, as well as the Support and Scholarship Fund administered by Sveriges Ingenjörer (The Swedish Association of Graduate Engineers) and the Sigurd and Elsa Golje Memorial Foundation.

## Data and code availability

All data are presented in the figures of this paper, and can be made available upon request from the authors. No new code was generated in this study.

## Materials and Methods

### Human and animal ethics

This study was approved by the UTS Research Ethics Committees with approval numbers ETH22-7590 and ETH20-5073. All urine donors provided informed consent for participation in this study under the provided ethics approval above. To follow sustainable guidelines in ethical animal research to reduce animals in research (*i.e.*, the 3 Rs), we only used bladders from naïve mice that were part of control groups in other studies or used for animal training at the animal facility at the University of Technology Sydney.

### Bacterial growth

Bacterial strains included in this study were: *Escherichia coli* K-12 strain MG1655, and uropathogenic *E. coli* strain UTI89.

**Table 1.**
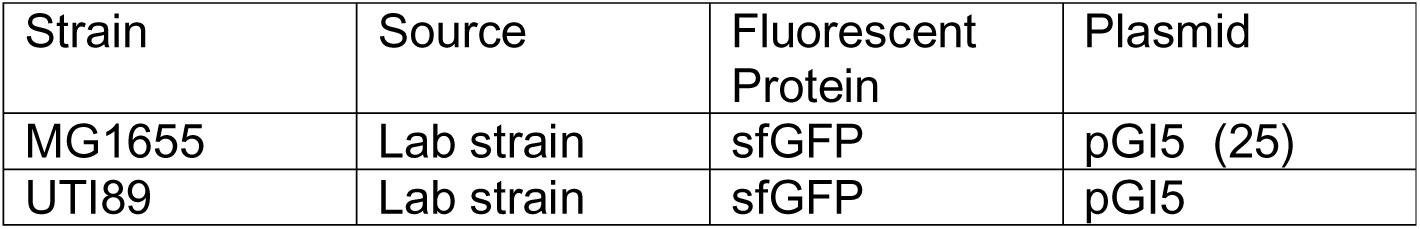
Strains and plasmids used in this study.

| Strain | Source | Fluorescent Protein | Plasmid |
| --- | --- | --- | --- |
| MG1655 | Lab strain | sfGFP | pGI5 (25) |
| UTI89 | Lab strain | sfGFP | pGI5 |

Transformed strains constitutively expressed green fluorescent protein (sfGFP) as freely diffusing molecules in the cytoplasm. Single colonies of the respective strains were grown overnight in 10 mL of Luria-Bertani broth supplemented with spectinomycin (100 µg mL^-1^) at 37 °C. The cultures were incubated statically to allow expression of type 1 fimbriae. To prepare for infections, overnight cultures of the respective strain were pelleted and resuspended in 1x PBS to an OD_600_ of 0.1.

### Urothelial cell maintenance

The immortalised bladder urothelial cell line PD07i (32) was maintained in EpiLife media supplemented with 1 % Human Keratinocyte Growth Supplement (HKGS) in 5 % CO_2_ at 37 °C. Cells were passaged twice per week using a standard trypsinisation protocol when approximately 80 % confluency was reached.

### Preparation of human urine

Human urine was collected from both male and female donors. Urine was never pooled. Samples were collected in the mornings and stored at 4 °C for 48 hours before downstream processing. Urine was centrifuged at 3900 rpm for 10 minutes, and the supernatant was passed through a 0.2 µm filter, aliquoted into 50 mL Falcon tubes, and stored at -20 °C until further use. To optimise the likelihood for filamentation, only urine with a urine specific gravity between 1.025 and 1.030 g mL^-1^ and a pH between 5 and 5.5 was used in this study (25, 44).

### Infection models

#### In vitro petri dish model

The *in vitro* petri dish model was described previously (37). In summary, 35 mm IBIDI glass bottom dishes with glass coverslip [#1.5] bottom (IBIDI #81218–200) were seeded with PD07i urothelial cells as per the manufacturer’s recommendations and allowed to grow until confluence. To assess E-cadherin and ZO-1 expression, cells were fixed with 4 % PFA in PBS for 10 minutes, followed by a permeabilising step with 0.1 % Triton X-100 for 10 minutes, and then blocked with 3 % BSA for at least 1 hour. The cells were then incubated with antibodies against E-cadherin (1:50, AB40772, Abcam) or ZO-1 (1:50, AB221547, Abcam) at 4 °C overnight. Following washes in PBS, a secondary anti-rabbit single-domain antibody conjugated to Atto488 (1:500, FluoTag®-X2 anti-Rabbit IgG, N2402-At488-S, NanoTag Biotechnologies, Germany) was added for 1 hour. After extensive washing in PBS, cell monolayers were covered in ProLong Gold (P36934, ThermoFisher) antifade mounting media prior to imaging.

For infection, dishes were incubated semi-statically (50 rpm orbital shaking) at 37 °C with 5 % CO_2_. Urothelial cells were challenged with *E. coli* strains at a multiplicity of infection (MOI) of 100 in PBS. The dishes were incubated for 15 minutes, before excess culture media was removed, 2 mL of fresh prewarmed EpiLife media supplemented with HKGS was added, and the dishes were allowed to incubate for another 6 hours. Media was replaced with 2 mL of EpiLife media (supplemented with HKGS) with gentamycin (50 µg mL^-1^), and incubated for 15 hours to kill extracellular bacteria. To observe bacterial invasion (after 6 hours) and the formation of IBCs (after 21 hours), PD07i cell membranes and nuclei were labelled with CellBrite Steady 405 and NucSpot Live 650 in EpiLife (1:100 dilution with the addition of the CellBrite Steady enhancer and Verapamil in accordance with the manufacturer’s recommendations) for 30 minutes. The cells were then washed three times with 1x PBS and replenished with 2 mL of fresh pre-warmed EpiLife media (supplemented with HKGS) to maintain cell viability throughout imaging.

#### In vitro adhesion assay

Bacterial adhesion assays were previously described (55) and used with modifications to perform similarly to the initial stages of the *in vitro* petri dish model. IBIDI 35 mm glass bottom dishes with glass coverslip [#1.5] bottom (IBIDI #81218–200) were seeded with PD07i bladder urothelial cells as per the manufacturer’s recommendations and allowed to grow until confluence. The urothelial cells were then challenged with *E. coli* strains at a MOI of 100 in PBS and incubated for 15 minutes at 37 °C with 5 % CO_2_. The cells were then washed three times with 1x PBS and lysed with 0.1 % Triton X-100 and 0.05 % Trypsin-EDTA at 37 °C with 5 % CO_2_ for 15 minutes. Adherent bacteria were then enumerated by plating serial dilutions on LB agar plates (supplemented with spectinomycin 100 µg mL^-1^) for CFU counting after overnight incubation at 37 °C.

#### In vitro microfluidics flow model

The *in vitro* flow model was previously described (25, 37) and used here with minor variations. On day one, PD07i bladder cells were seeded into µ-Slide I^0.2^ Luer (IBIDI #80166, total channel volume 50 µL) flow channels. The channels were incubated overnight to allow the cells to grow to confluence. On day two, the channels were connected to a New Era pump system to allow continuous media flow. To initiate infection, *E. coli* strains were resuspended in PBS to an OD_600_ of 0.2 and were then connected to the flow channel directly and flowed at 50 µL min^-1^ for 15 minutes. Fresh EpiLife supplemented with HKGS was then reconnected to the system and flowed at an initial rate of 100 µL min^-1^ for 10 minutes to flush unbound bacteria, then 15 µL min^-1^ for 6 hours. Following this, EpiLife supplemented with HKGS and 50 µg mL^-1^ gentamycin was flowed at 15 µL min^-1^ for 18-20 hours, to kill and remove extracellular bacteria. To induce infection-related filaments, the growth media was exchanged for human urine at a flow rate of 15 µL min^-1^ for an additional 18-20 hours. For infections without urine, EpiLife supplemented with HKGS was exchanged into the system at a flow rate of 15 µL min^-1^ for an additional 18-20 hours. Bacteria were then harvested from the back outlet of the flow channels and resuspended in PBS for imaging.

For urine semi-static (50 rpm orbital shaking) dish infections, a combination of the *in vitro* dish model and the flow model timepoints was utilised. On day one, PD07i bladder cells were seeded into 35 mm IBIDI glass bottom dishes with glass coverslip [#1.5] bottom (IBIDI #81218–200) and were incubated overnight to grow to confluence. To initiate infections on day two, *E. coli* strains were resuspended in PBS to an OD_600_ of 0.2. The bacteria were then directly added to the dish and allowed to incubate at 37 °C for 15 minutes. The excess PBS was then removed, and 2 mL of prewarmed EpiLife media (supplemented with HKGS) was added and incubated for a further 6 hours. To kill extracellular bacteria, EpiLife (supplemented with HKGS) with gentamycin (50 µg mL^-1^) was exchanged into the dish and incubated for 18-20 hours. To induce infection-related filamentation, EpiLife media was removed and exchanged with human urine and allowed to incubate for 18-20 hours. Bacteria were then harvested from the supernatant of the dish and resuspended in PBS for imaging.

#### Ex vivo mouse urothelial sheet model

The *ex vivo* model was previously described (45). Briefly, bladders used in this study were extracted from donated 6–8-week-old C57BL/6J mice. Harvested bladders were stored in cold PBS until infection (maximum storage time before infection <24h, as this does not significantly affect infection dynamics (45)). Bladders were dissected into quarters, and the urothelial layer was separated from the muscle layer using forceps underneath a dissecting microscope and placed into the first column of a U-bottom 96-well plate containing PBS. To initiate infection, bacterial cultures were resuspended to an OD_600_ of 0.1 in PBS and pipetted into the second column. Using forceps, the bladder sheets were moved from the first column to the second, and the plate was incubated for 4 hours at 37 °C with 5 % CO_2_. To kill unbound, extracellular bacteria, the sheets were moved to column 3 containing PBS supplemented with gentamycin 100 µg mL^-1^ and incubated for 1 hour. Urothelial sheets were then washed with 1x PBS in columns 4 and 5 and then incubated overnight in PBS in column 6 at 37 °C with 5 % CO_2_. To observe IBC formation, urothelial sheets were labelled with either NucRed Live 647 or NucSpot Live 650 in PBS (1:100 final concentration as recommended by the manufacturer) for 15 minutes. Sheets were washed three times with 1x PBS and carefully spread onto an agarose pad (2 % W/V) for imaging.

To induce infection-related filamentation, infected sheets were placed in columns 2 to 8 in the following row of the U-bottom 96-well plate containing human urine (pH between 5.12 and 5.5 and urine specific gravity of at least 1.025 g mL^-1^). To supply the sheets with fresh urine, the urothelial sheets were incubated for 1 hour before being moved to the next well for the first 6 hours (*e.g.* initial incubation in column 2, then moved to column 3 for another hour of incubation, etc. This was done to expose the sheet to fresh urine for the first part of the infection. After the sixth change, the urothelial sheets were incubated overnight in urine. After overnight incubation, both the urothelial sheets and the supernatant were collected and processed for imaging.

### Imaging

For the live cell *in vitro* and *ex vivo* infections, imaging bias was minimised by selecting regions of interest (ROIs) based on the brightfield channel (*in vitro*) or nucleus staining (*ex vivo* mouse sheets) of a Leica Stellaris 8 confocal microscope mounted with a 63x 1.40 NA oil-immersion objective and an environmental chamber set at 37 °C. Z-stacks were then acquired on the selected ROIs to validate that the bacteria were localised intracellularly in the host cells. Fluorophores were excited by a white laser at optimised wavelengths for each fluorophore, and the emission was collected using pre-set, system-optimised detector ranges for AlexFluor405, EGFP, and AlexaFluor647, depending on the experiment, to minimise channel crosstalk. Image size was either 512 x 512 or 1024 x 1024 pixels. The pinhole was set to 1 AU, and Z-stacks were collected with a software optimised step length of 500 nm (42–162 images per stack, depending on the thickness of the bladder cell or sheet in question).

3D reconstruction and deconvolution of Z-stacks were performed on the Leica LAS X software, visualised by Imaris software (10.0.1), and further analysed in FIJI (ImageJ). Monolayers of fixed immuno-stained cells were imaged using a Nikon N-STORM Ti2-E NIS v.5.30. fluorescence microscope operated in TIRF mode (with HILO illumination set to a few degrees above the critical angle), mounted with a 100x 1.49 NA oil immersion objective. Fluorophores were excited using 488 nm and 647 nm laser lines, and fluorescence emission was detected using a sCMOS Flash 4.0 v3 (Hamamatsu) camera after emission was filtered through standard filter cubes for FITC and ‘Normal STORM(647)’. Image acquisition times were set between 50 and 150 ms, and the image size was 2048 x 2048, with a pixel size of 65 nm.

To assess the viability and reversal of the filaments from the flow- and *ex vivo* models, live-cell epifluorescence and bright-field time-lapse microscopy of filaments reverting back to rod-shaped cells was conducted using a Nikon N-STORM Ti2-E fluorescence microscope, mounted with a 100x 1.49 NA oil immersion objective, and an environmental chamber set at 37 °C (Okolab cage incubator). ∼ 2 µL of samples were placed on pre-made 1.5 % agarose pads in LB media in 65 µm gene frames (Thermo Scientific, AB0577). Intracellular GFP was excited using a Lumencore Specta II module and detected using a FITC filter cube. Images were acquired every 10 minutes using a sCMOS Flash 4.0 v3 (Hamamatsu) camera.

### Immunoblotting

Bacterial cell extracts from a volume corresponding to 0.1 OD_600_ units were collected for all strains. The extracts were suspended in loading buffer and resolved by SDS-PAGE. Proteins were transferred to nitrocellulose membranes using a rapid Transfer-Blot apparatus (Bio-Rad). The membranes were blocked in 5 % (w/v) milk and probed with antibodies for FimH 1:10000 in TBST (Genescript, Singapore) overnight at 4 °C. The following day, membranes were washed 3 times in TBST and probed with secondary HRP-antibodies (1:10000, goat anti-rabbit, Bio-Rad, #1706515) for 1 hour before being developed and imaged.

### Quantification and statistical analysis

To assess invasion frequencies of the bladder cells, microscopy images and deconvolved Z-stacks were analysed using the Cell Counter plugin in FIJI (ImageJ). Due to the complexity of the filament samples (cells often overlapping and crossing), length analysis was conducted by manually tracing a line along the midline of the cell in FIJI (ImageJ) using the segmented line tool. Statistical analysis was performed in GraphPad Prism 10.2.3 and OriginPro 2021 V9.8.0.200 (academic). To compare the means of two independent groups, an unpaired T-test was used. A one-way ANOVA was used to compare the means of three or more independent groups. Levels of significance were indicated as: ns, not significant; *, *P* < 0.05; **, *P* < 0.01, ***, *P* < 0.001.

