## Supplementart info for "Invasion and intracellular proliferation of non-pathogenic *E. coli* in model infection systems"

Supplementary figures

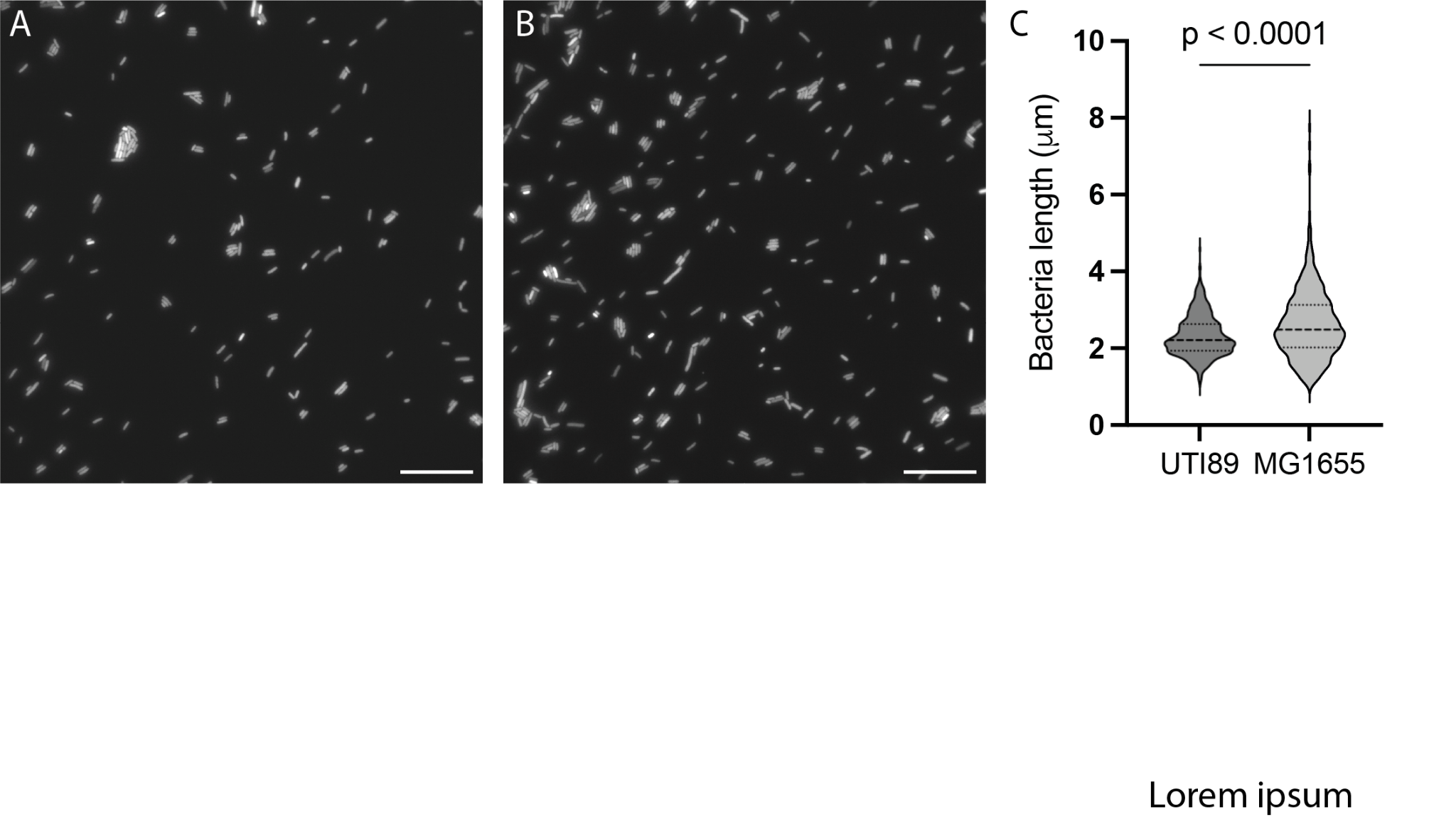

Supplementary Figure S1. Growing bacteria in urine overnight (37 °C, static) does not induce infection-related filamentation. Representative images of **A**, UTI89 and **B**, MG1655. Scale bar 20 µm. Note same batch of urine was used as for the overnight *ex vivo* mouse bladder sheet infections. **C**, Graph shows mean lengths of $\bar{L}$_UTI89_ = 2.32 µm and $\bar{L}$_MG1655_ = 2.63 µm (n > 450 cells per strain (n = 3))
